# FlowPacker: Protein side-chain packing with torsional flow matching

**DOI:** 10.1101/2024.07.05.602280

**Authors:** Jin Sub Lee, Philip M. Kim

## Abstract

Accurate prediction of protein side-chain conformations is necessary to understand protein folding, proteinprotein interactions and facilitate de novo protein design. Here we apply torsional flow matching and equivariant graph attention to develop FlowPacker, a fast and performant model to predict protein sidechain conformations conditioned on the protein sequence and backbone. We show that FlowPacker outperforms previous state-of-the-art baselines across most metrics with improved runtime. We further show that FlowPacker can be used to inpaint missing side-chain coordinates and also for multimeric targets, and exhibits strong performance on a test set of antibody-antigen complexes. Code is available at https://gitlab.com/mjslee0921/flowpacker.

## 1 Introduction

A protein’s three dimensional structure, determined by its primary amino acid sequence, is the main determinant of its function. The side-chain atoms heavily influences its folding and interaction with other proteins/ligands through various interatomic interactions and potentials. Therefore, to fully understand protein folding and identify protein-protein interactions, accurate models of protein side-chain packing must be developed.

Recent advances in artificial intelligence has made astounding progress in protein structure prediction [1, 2] and design [3, 4, 5]. Here we focus on the problem of side-chain packing, which seeks to predict the side-chain conformations given the amino acid sequence and backbone coordinates of a protein structure. Many prior efforts rely on physics-based modeling, which uses empirical scoring functions [6], discrete rotamer libraries [7], and/or MCMC-based sampling to identify plausible rotamers. However, these methods are often ineffective due to inefficient search algorithms and the reliance of often inaccurate scoring functions that converge to suboptimal local minima. Recent deep learning-based methods have shown significant improvements in runtime and efficacy to physics-based modeling in side-chain packing [8, 9, 10, 11], the most notable of which is DiffPack [10], a torsional diffusion model that autoregressively generates the four *χ* torsion angles that constitute the only degrees of freedom for side-chain conformations. DiffPack presents a number of innovations, such as autoregressive generation and confidence sampling, to attain state-of-the-art performance in protein side-chain packing. In this work, we approach the side-chain packing problem using flow matching and equivariant graph attention networks to attain state-of-the-art performance.

Flow matching [12] is a novel generative modeling paradigm that allows the training of continuous normalizing flows (CNFs) in a simulation-free manner, and have shown stronger performance, faster training convergence, and faster inference than standard diffusion models. In FlowPacker, we replace the torsional diffusion framework with torsional flow matching, derived from flow matching on Riemannian manifolds [13]. We also apply state-of-the-art equivariant graph attention models - namely, EquiformerV2 [14] - which has been shown to improve performance over invariant message passing networks due to increased expressivity and parameter efficiency.

We observe FlowPacker outperforms other side-chain packing baselines across most evaluated metrics while being considerably faster. We show that FlowPacker can be used for partial inpainting of side-chain conformations, a feature not readily available with other packing methods. We also show that FlowPacker can be extended to multimeric complexes and specifically with antibody-antigen complexes and show significant performance improvements in CDRH3 and full variable chain (Fv) side-chain packing. Therefore, it can be easily incorporated with existing backbone generative models and sequence design tools to generate accurate full-atom structures.

## 2 Background

### Preliminaries

Proteins are macromolecules that adopt three-dimensional conformations based on its amino acid identity and resulting covalent and non-covalent interactions between atoms. The atoms that constitute a full-atom protein structure are divided into backbone atoms, which are the N, Ca, C, and O atoms that constitute the peptide backbone, and side-chain atoms, which are dependent on the amino acid identity and can contain up to 9 heavy atoms in total. Side-chain flexibility is usually limited to four degrees of freedom defined by the four *χ* torsion angles whose empirical distributions are highly constrained based on amino acid type and neighboring atomic forces. The problem of side-chain packing, therefore, can be reduced to finding *χ*_1…4_ *∈* [0, 2*π*) conditioned on the protein sequence *s ∈ {*0, …, 19*}* and backbone coordinates *X*_*bb*_ *∈ R*^3^.

### Related Work

Side-chain packing methods has traditionally relied on energy-based sampling such as Rosetta [6] or SCWRL [15], but has largely been replaced with the rise of performant deep learning methods. Notably, AttnPacker [9] is a model based on AlphaFold2 [1] that uses sparse triangular attention and invariant point attention to directly predict all-atom coordinates. DiffPack [10] proposes a diffusion-based approach that applies a relational graph convolution network on atom-level graphs, with several innovations such as autoregressive sampling to attain state-of-the-art performance. SidechainDiff [16] presents a sidechain packing diffusion model for the prediction of mutational effects using transfer learning. More recently, two peptide-specific torsional flow matching models have been published, where PPFlow [17] uses torsional flow matching on both backbone and side-chain torsion angles for peptide design, while PepFlow [18] uses a multi-modal approach to full-atom peptide design against protein pockets, using *S*(*E*)3 flow matching for backbone generation, simplex flow for amino acid sequences, and torsional flow matching for side-chain *χ* angles.

## 3 Methods

A schematic of FlowPacker is provided in Figure 1. Here we provide a brief overview of the theoretical background and model details.

### 3.1 Torsional Flow Matching

Continuous normalizing flows (CNFs) are a class of generative models that uses a learned vector field to transform a simple prior to the desired data distribution. However, the maximum likelihood simulation-based training of CNFs have limited its scalability and application to complex datasets. Recently, flow matching was proposed to train continuous normalizing flows (CNFs) in a simulation-free manner. Lipman et. al. [12] show that CNFs can be trained by regressing on a conditional probability path *p*_*t*_(*x*|*x*_1_) to learn the unconditional path *p*_*t*_(*x*) that transforms a prior density *p*(0) to the data distribution *p*(1). Chen et al. [13] extends the flow matching framework to Riemannian manifolds including the high-dimensional torus, which is used in this work to define a torsional flow matching framework for side-chain conformation generation and briefly discussed below.

In flow matching, we desire to learn the time-conditioned vector field *v*_*t*_(*x*) with *t ∈* [0, 1] that transforms a simple prior distribution *p*(0) to the data distribution *p*(1), where the learned vector field can then be used to integrate a prior sample from *t* = 0 to *t* = 1 for generative modeling. However, the probability path *p*_*t*_(*x*) is intractable to compute. Lipman et. al. [12] show that we can use the *conditional vector field v*_*t*_(*x*|*x*_1_) that generates *conditional probability paths p*_*t*_(*x*|*x*_1_) as regression targets, which converges to the same optima as the unconditional vector field *v*_*t*_(*x*) that produces the unconditional probability path *p*_*t*_(*x*). Given a conditional flow *ψ*_*t*_(*x*_0_|*x*_1_) that transforms the prior distribution *p*_0_(*x*|*x*_1_) to *p*_*t*_(*x*|*x*_1_), we observe that:

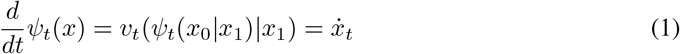

which defines a conditional vector field that can serve as the regression target for simulation-free training, resulting in the conditional flow matching (CFM) loss below:

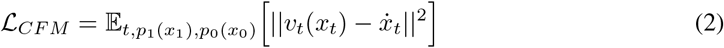

Chen and Lipman [13] introduce Riemannian flow matching, which extends flow matching to manifolds and general geometries. They show that the conditional flows *ψ*_*t*_(*x*|*x*_1_) can be constructed on simple manifolds using a linear scheduler *κ*(*t*) = 1 *‐ t* with the geodesic distance as the *premetric* that concentrates all mass at *x*_1_ for *t* = 1. The geodesic distance for simple manifolds, such as the high-dimensional torus considered in this work, can be computed in closed form with the logarithmic map between two points on the manifold, and mapped back to the manifold using the exponential map. Thus, the conditional flow is defined as:

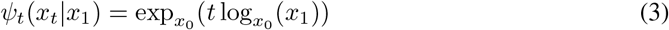

where the exponential and logarithmic map for high-dimensional tori are defined as:

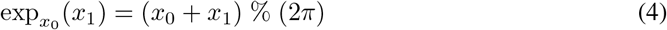

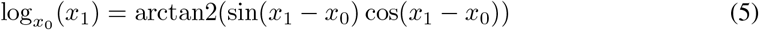

This corresponds to a flow along the geodesic distance between *x*_1_ and *x*_0_ that varies linearly with time. This can be used to interpolate *x*_*t*_ in (2) to define the torsional flow matching loss used in this work.

### 3.2 Equivariant graph attention using Equiformerv2

Equivariant neural networks have been shown to increase model performance and efficiency by directly integrating the symmetries of the data into the model architecture, without the need for data augmentation nor additional parameters [19]^1^. Equiformer [21] is a graph attention network that operates on equivariant irreducible representation (irreps) features and transfers information between various type-*L* vectors using tensor product operations, where different type-*L* vectors can be combined through the use of Clebsch-Gordan coefficients. Equiformer uses depthwise tensor product (DTP) blocks - where one type-*L* vector in the output irreps is only dependent on one type-*L*’ vector in the input irreps - to reduce computational complexity of tensor product, but is still practically restricted to max *L* = 3 vectors due to efficiency. For a more detailed discussion on tensor product operations, we refer readers to [22] and [23].

Equiformerv2 [14] facilitates the scaling of the model to higher-degree features by replacing *SO*(3) convolutions with eSCN convolutions [24], which intuitively aligns the relative edge direction to one axis and therefore greatly sparsifies the Clebsch-Gordon coefficient matrix and simplifies to *SO*(2) convolutions, reducing computational complexity and allowing scaling to higher-degree tensors up to *L* = 6 or *L* = 8 for increased model expressivity. While the task of side-chain packing does not necessitate the use of equivariant networks since torsion angles are invariant features, in preliminary studies we observed stronger empirical performance of Equiformerv2 over invariant and equivariant baseline architectures.

### 3.3 FlowPacker specifications

#### Dataset curation

We train FlowPacker on two datasets: 1. BC40 dataset (release date 2020-07-28, available at https://drug.ai.tencent.com/protein/bc40/download.html), which are representative PDB structures clustered by MMseqs2 [25] at 40% sequence identity, and is used by previous side-chain packing models (DiffPack [10] and AttnPacker [9]) for model training, and 2. a monomer dataset constructed from a PDB snapshot (date 2023-07-28) clustered at 40% sequence identity (hereafter referred to as PDB-S40). All results are based on the model trained using PDB-S40 unless otherwise noted. We use the CASP13, 14, and 15 targets for testing (available at https://predictioncenter.org/download_area/). We reduce data redundancy in the training set by removing any structures/clusters with >40% sequence similarity to any of the CASP13/14/15 targets using MMseqs2’s `easy-search` workflow. We discard structures of which more than 25% of residues are unknown, and remove residues with missing backbone coordinates or overlapping alpha carbon positions. We also filter for proteins with at least 40 residues. This results in 36,451 training examples for the BC40 dataset and 23,191 clusters containing 309,467 structures for the PDB-S40 dataset. The test set consists of 99 structures, with 20, 34, and 45 targets in CASP13, 14, and 15, respectively.

**Figure 1:**
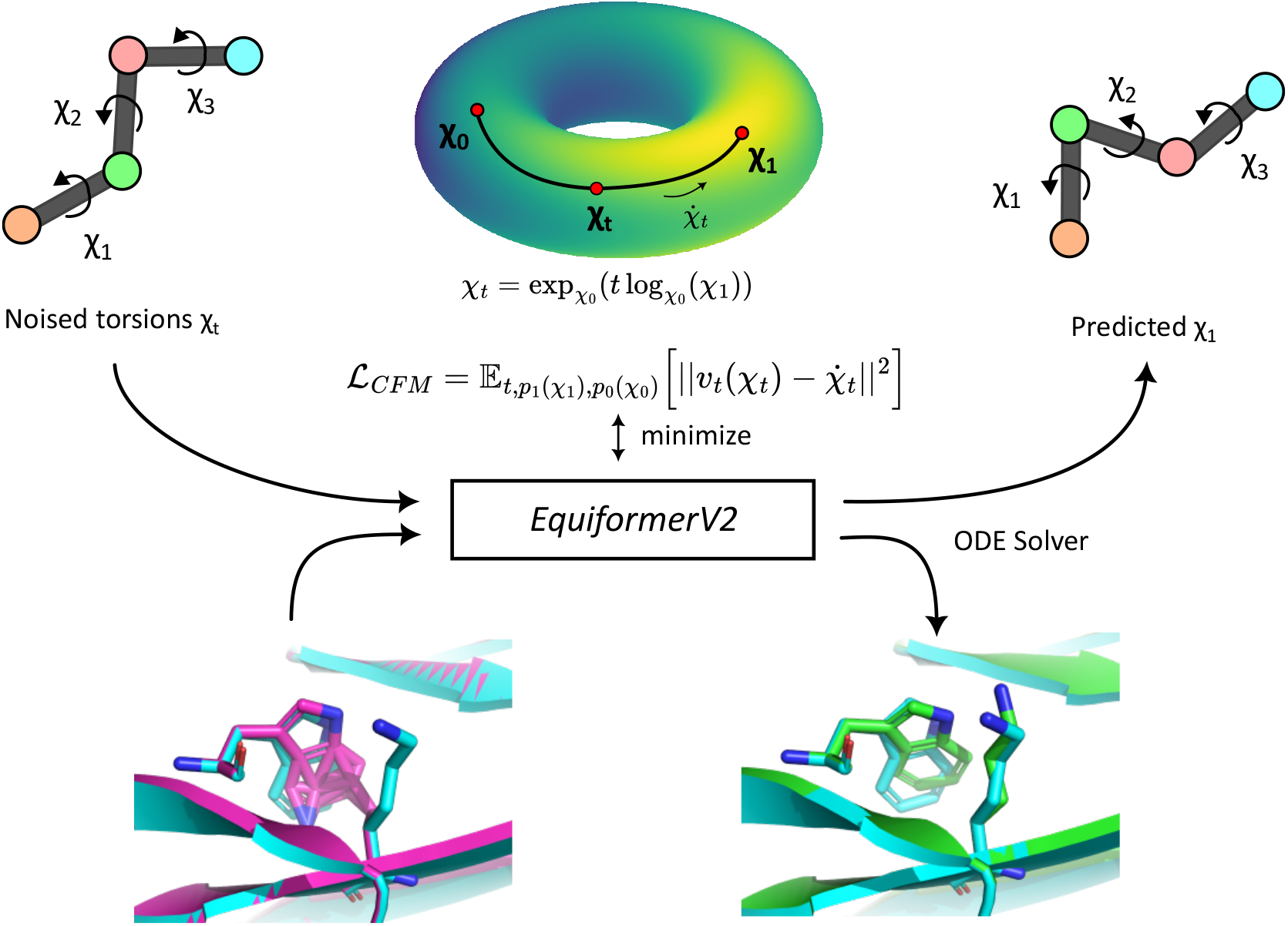
FlowPacker Overview. FlowPacker is an equivariant graph attention network that generates side-chain conformations of a given protein structure and sequence using torsional flow matching. The model predicts the vector field of the conditional flow along the hypertorus between a prior angle *χ*_0_ and the ground-truth angle *χ*_1_, which is used with an ODE Solver (ex. Euler’s method) to generate a sample from the data distribution. Note we show a ‘virtual’ side-chain conformation with three *χ* angles, but each amino acid can contain up to 4 *χ* angles that affect the positions of up to 9 atoms.

#### Model architecture

FlowPacker uses the Equiformerv2 model architecture as described previously. We use *L*_*max*_ = 3, channel dimension of 256, and 4 blocks, resulting in a total of 18.0M trainable parameters. Interestingly, we did not observe a change in performance from scaling *L*_*max*_ up to 6, suggesting that higher-order features are not informative for side-chain packing.

#### Loss functions

The model is trained to predict the conditional vector field as defined in Section 3.1:

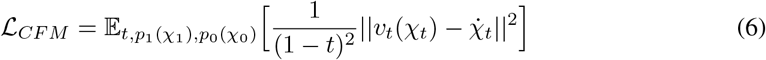

where *v*_*t*_(*χ*_*t*_) corresponds to the model output, 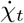 is the exact conditional vector field efficiently computed with autograd, and 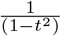 is a weighting factor.

**Table 1:**
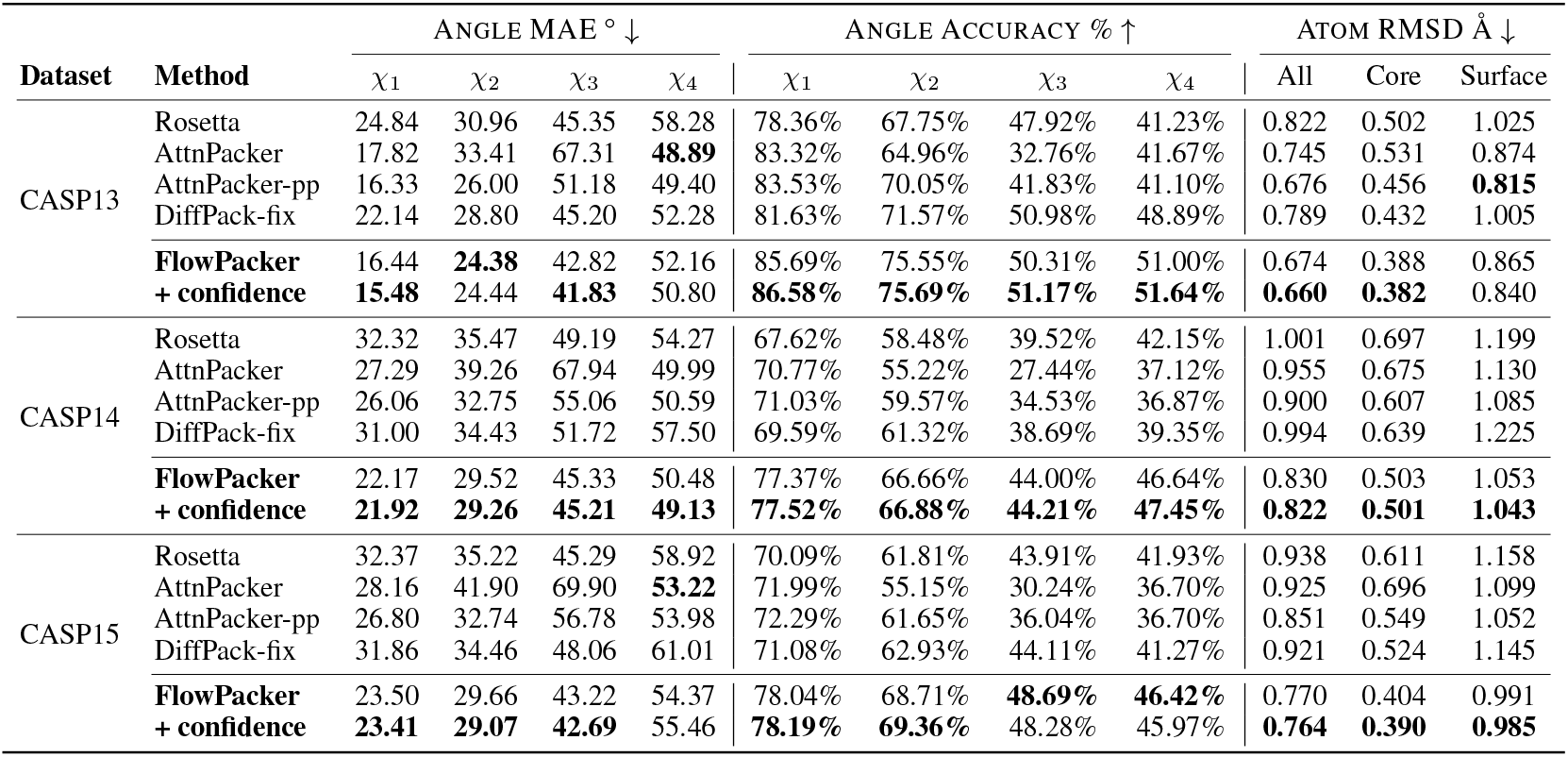
Performance evaluation of FlowPacker and other methods on CASP targets. The top performing model per test dataset is highlighted in **bold**. DIFFPACK-FIX shows results for DiffPack using structures with idealized bond lengths and angles as input. We generate 4 samples for FlowPacker (displaying average metrics when confidence model is not used) and use default settings for DiffPack (4 samples with confidence model selection). AttnPacker-pp corresponds to samples with post-processing applied as denoted in [9].

We experimented with multiple variations of the loss function including using the vector field approximation log_*χt*_ (*χ*_1_)*/*(1 *‐ t*), which showed negligible differences in training dynamics and performance. We also tried using a reparameterized direct *χ*_1_ prediction as with rotation matrices in FrameFlow [4], which can be used to approximate the vector field during sampling. However, this caused unstable training dynamics, most likely due to the degeneracy of *χ* angles.

#### Handling symmetry issues

Certain *χ* angles exhibit *π*-symmetry that may adversely affect model efficiency. We handle these cases by taking mod *π* for *π*-symmetric *χ* angles and modifying equation (4) to:

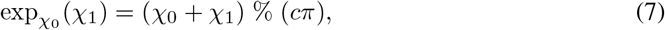

where *c* = 1 for *π*-symmetric *χ* angles and *c* = 2 otherwise, restricting *π*-symmetric angles to [0, *π*).

#### Training details

In FlowPacker, we use an residue-level graph, where each residue represents a node with the position given by its idealized C*β* atom. At each training step, we sample timesteps *t* and generate noised torsions *χ*_*t*_, which are used to reconstruct noised atomic coordinates *X*_*t*_ using idealized coordinates. The node features consist of the amino acid identity, backbone torsion angles, and sinusoidal embeddings of the current timestep, while the edge features are relative positional encoding clamped at [*‐*32, 32], and the Euclidean distance between all idealized coordinates (using the atom14 representation) between two connected residues. The edges are defined by a *k*-nn graph with *k* = 30. The model is trained on 4 40GB NVIDIA A100s, effective batch size of 16, and context length of 512 residues (cropped when length is greater than 512) for 300 epochs, or approximately 6 days. We use the AdamW optimizer with learning rate 1.0*e*^*‐*4^ and gradient clipping at a norm value of 1.0.

#### Inference strategies

Though the model was trained using a conditional flow with a linear scheduler *κ*(*t*) = 1 *‐ t*, we observe a similar phenomenon as described in FrameFlow where the generated samples are poor using the same linear schedule during sampling. Instead, we adopt the exponential schedule *v*_*t*_ = *c*(log_*xt*_ (*x*_1_)), and empirically find that c = 5 works well. We use a uniform distribution on *SO*(2) as the base distribution and Euler solver with t = 10 for all sampling steps, as we observed no significant increase in performance with additional timesteps. We experimented with using higher- order solvers such as Heun’s method and RK45 solvers, but did not observe a noticeable performance improvement at the cost of increased runtime. We use exponential moving average with decay of 0.999 at every training iteration.

**Figure 2:**
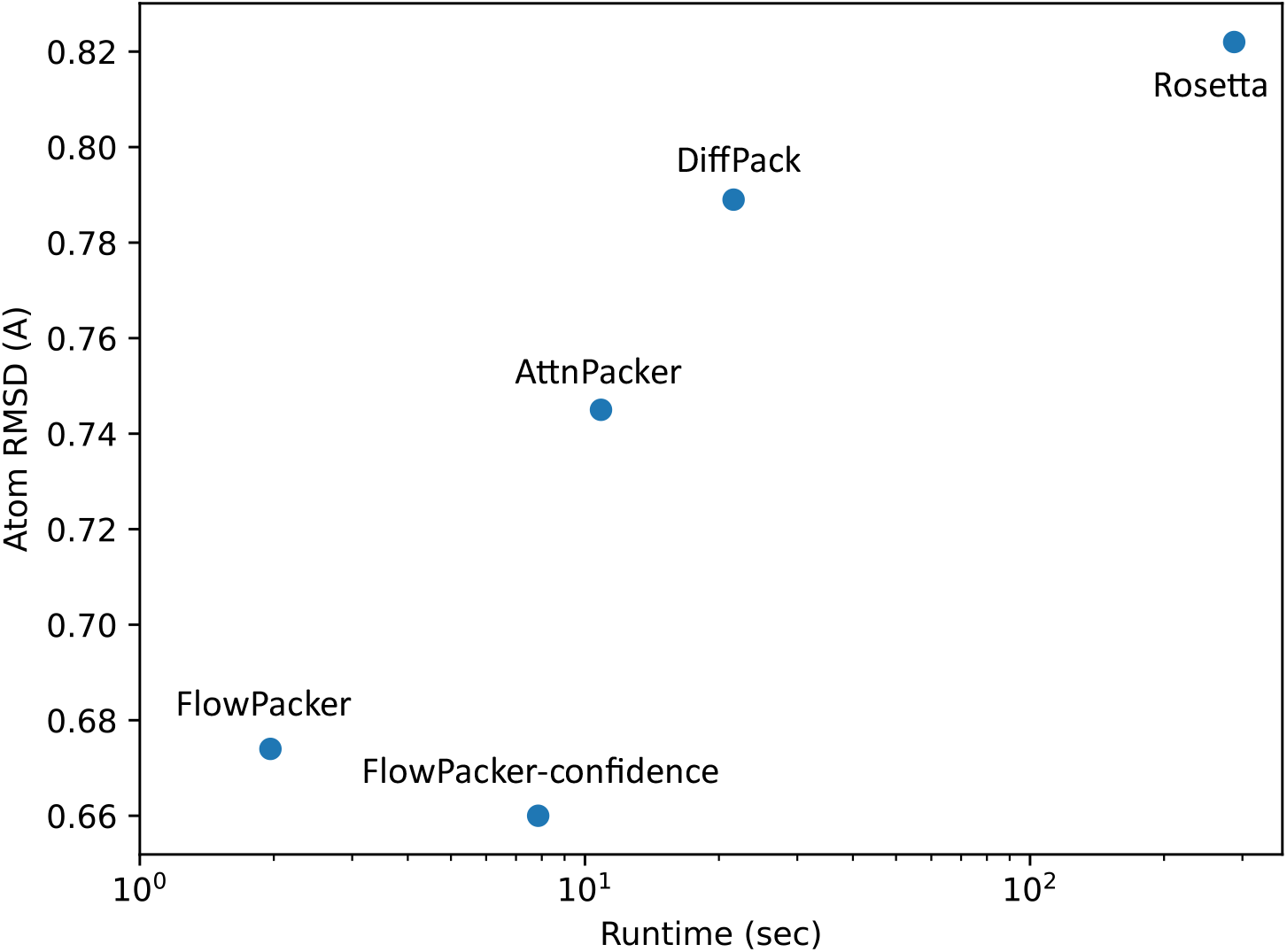
Runtime and performance analysis of various methods. We use the CASP13 dataset to analyze the total runtime (averaged per protein) and Atom RMSD across the tested methods. FlowPacker exhibits the best RMSD-runtime tradeoff, while FlowPacker with the confidence model generates conformations with the lowest RMSD. We report the metrics of DIFFPACK-FIX in Table 1 here, and all runtime metrics are averaged over three independent seeds for each method. All models were tested on AMD Ryzen 7 5800 (Rosetta) or NVIDIA RTX 3060 (others). Note that runtime is plotted in log scale.

#### Confidence model

The confidence model is trained in a similar manner to the main model, except we regress on the residue-level sidechain RMSDs rather than *χ*-level vector fields. We identify the best sample by simply taking the mean predicted RMSD across all residues and selecting the sample with the lowest predicted RMSD. Unless otherwise noted, we generate 4 samples per test case and select the highest confidence sample. The confidence model trained in this work contains 2.2M trainable parameters with max *l* = 2 and hidden dimension of 64 - we did not perform extensive hyperparameter tuning on this module.

## 4 Results

We compare FlowPacker to physics-based (Rosetta [6]) and deep learning-based (AttnPacker [9] and DiffPack [10]) methods across three test datasets - CASP13, CASP14, and CASP15 - in Table 1. As with previous studies [9, 10], we report three metrics: 1. ANGLE MAE corresponds to the angular mean absolute error, which returns min(*χ, χ* mod 2*π*), 2. ANGLE ACCURACY refers to the percentage of angles within 20^*°*^ of the ground-truth, and 3. ATOM RMSD refers to the average root-mean squared deviation of side-chain atoms per residue, where core residues are defined as residues with at least 20 *Cβ* atoms within 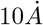 , and surface residues are those with at most 15 *Cβ* Within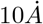.

**Table 2:** Performance on CASP15 targets with varying levels of masking. We mask varying proportions of residues randomly and calculate Angle MAE and Atom RMSD on the masked residues.

| Masking | ANGLE MAE ° ↓ |  |  |  | RMSD Å ↓ |
| --- | --- | --- | --- | --- | --- |
| | $\chi_1$ | $\chi_2$ | $\chi_3$ | $\chi_4$ | All |
| 5% | 21.51 | 29.07 | 50.33 | 54.51 | 0.727 |
| 10% | 21.85 | 27.62 | 44.46 | 52.39 | 0.728 |
| 25% | 22.37 | 28.05 | 45.03 | 53.34 | 0.737 |
| 50% | 23.12 | 29.03 | 43.40 | 53.63 | 0.756 |
| 75% | 23.30 | 29.60 | 43.30 | 54.05 | 0.765 |
| 100% | 23.50 | 29.66 | 43.22 | 54.37 | 0.770 |

During testing, we observed a data leakage issue with DiffPack, where the generation quality was dependent on the input side-chain coordinates, everything else held constant. We noticed that during training and sampling, DiffPack rotates the ground-truth side-chain coordinates given *χ*_*t*_, maintaining the ground-truth bond angles and bond lengths. Moreover, we discovered that if we idealize the side-chain atoms (as in AlphaFold2 [1] and FlowPacker) and provide the idealized structures (note we still use the non-idealized ground-truth backbone coordinates) to DiffPack, the generation quality is significantly worse (see DIFFPACK-FIX in Table 1). We hypothesize that although the *χ*_1…4_ angles are the primary degrees of freedom in side-chain packing, the atomic coordinates do exhibit minor deviations from various interatomic forces and potentials [26] that may provide clues into the ground-truth side-chain conformation. Moreover, we observed that using ground-truth bond angles and lengths provides a >0.25A RMSD advantage over using idealized ones in terms of side-chain RMSD. For further discussion, please refer to Supplementary Information 1.1. Since side-chain packing methods should require no *a priori* knowledge of ground-truth side-chain positions, we exclude DiffPack from ranking across metrics. We note that the numbers denoted under DIFFPACK-FIX may not accurately reflect the true performance of DiffPack since the model is not explicitly trained with idealized coordinates - however, this requires further investigation that is outside of the scope of this work.

We observe that FlowPacker outperforms all baselines across most metrics, suggesting the superiority of flow matching and equivariant networks over diffusion and invariant models, respectively. Using a confidence model to select the lowest predicted RMSD sample marginally improved performance across all test datasets. However, the < 0.1 RMSD decrease suggests that for high-throughput screening, the confidence model may not be necessary since it requires multiple generations (4 used here), each with an extra forward pass through the confidence model that results in a > 4X increase in runtime. We observe that *χ*_1_ accuracy is generally higher than the subsequent *χ* angles, as seen with previous methods, due to the lever effect where the errors accumulate through each *χ* angle and the increased flexibility of side-chain positions at longer side-chains (ex. arginines and leucines). We also analyze runtime and performance across all methods in Figure 2, and observe that FlowPacker exhibits the best runtime and performance of all tested baselines. As shown in Supplementary Figure 2, FlowPacker can recapitulate the *χ* distributions with decent accuracy, including *π*-symmetric ones, which lie in the interval [0, 180] due to the parameterization in Equation 7. An analysis of residue-level atom RMSDs for each unique amino acid reveals that longer and aromatic sidechains are often harder to predict accurately over shorter ones Supplementary Figure 3. We additionally observe that the increased training data available in the PDB-S40 dataset consistently improves performance across most metrics over the widely used BC40 dataset Supplementary Table 2, suggesting the importance of careful training data curation.

To test FlowPacker’s ability to inpaint missing side-chain coordinates, we mask 5-75% randomly selected residues on the CASP15 test set and report the metrics in Table 2. As expected, we observe lower RMSDs as we provide more structural context, suggesting the utility of FlowPacker for conditional design. We expect this to be useful in common protein design cases such as motif-scaffolding or interface design, where protein backbone generative models design the backbone coordinates, and FlowPacker can be used in conjunction with sequence design tools to generate full-atom coordinates.

**Table 3:** Test case on side-chain packing on antibody-antigen complexes.

| Target | Method | ANGLE MAE $^{\circ}$ ↓ | | | | RMSD $\text{\AA}$ ↓ |
| --- | --- | --- | --- | --- | --- | --- |
| | | $\chi_1$ | $\chi_2$ | $\chi_3$ | $\chi_4$ | All |
| CDRH3 | Rosetta | 28.05 | 26.04 | **38.16** | 58.64 | 0.886 |
|  | **FlowPacker** | **20.85** | **25.06** | 42.16 | **51.67** | **0.671** |
| Full Fv | Rosetta | 30.28 | 30.72 | 48.50 | 54.83 | 0.790 |
|  | **FlowPacker** | **22.79** | **24.32** | **41.71** | **51.03** | **0.649** |

Although FlowPacker was specifically trained on single-chain proteins, we tested whether the model can be used to generate side-chain conformations of antigen-antibody complexes. We extract antibody-antigen complexes from SabDab [27] and filter for structures released after 2021-01-01. We also remove complexes with at least 40% antigen sequence similarity with MMseqs [25] to any complex structure published prior to 2021-01-01 to remove redundancy, and crop antigens to 256 residues proximal to the antibody. This results in 104 clusters, from which we select a representative member (provided by MMseqs) from each cluster for testing. For inference on multi-chain proteins, we simply fix the relative positional encodings for edges between residues of different chains to 32 and all other input features are kept the same as in single-chain inference. In Table 3, we report the metrics for two common design tasks: CDRH3 design and full Fv design. For all cases, we provide the antigen structure as conditional inputs and pack the sidechains of the CDRH3 or full Fv region (or heavy chain only if light chain data is not available). We only use Rosetta as the baseline, as the other baseline methods are not adapted for multimeric inputs. FlowPacker outperforms Rosetta in both tasks and across most metrics despite not being explicitly trained on multimeric complexes. Therefore, FlowPacker can be used to generate more accurate full-atom structures of both monomeric and multimeric proteins, not limited to antibody-antigen complexes.

## 5 Discussion

In this work, we present FlowPacker, a torsional flow matching model that exhibits superior performance and runtime efficiency over other baselines in side-chain packing. We show that FlowPacker can be used to inpaint partial residues and multi-chain inference, showcasing a test case with antibody-antigen CDR side-chain packing, outperforming other methods in angle accuracy and atom RMSD.

We see various promising avenues for future work, such as improved prediction of mutational effects using unsupervised [28] or supervised [16] learning, or alignment of generative models using preference data [29, 30] such as empirical force fields to increase biophysical plausibility. We also believe that FlowPacker’s performance can be improved - we did not test autoregressive sampling as in DiffPack, which may help performance. We show that the confidence model modestly improves performance, but there are other applications such as resampling of low-confidence conformations and uncertainty analysis as with pLDDT in AlphaFold that may be further explored. We also encourage the exploration of novel representations of side-chain conformations, as explicit representation in 3D space [31, 2] may outperform implicit representations since the atom RMSD using a *χ* angle parameterization is often largely dependent on the accuracy of the *χ*_1_ prediction. Finally, we envision accurate side-chain packing models to be increasingly necessary with the development of more powerful protein backbone generative models to provide a pipeline for end-to-end full-atom protein design.

## 6 Acknowledgements

We thank the many open-source codebases that this work was based on, including PyTorch, OpenFold, and e3nn. We also would like to thank Digital Research Alliance of Canada for computing resources.

## Supplementary Information

### 1.1 Performance issues with DiffPack

The discrepancy in performance of DiffPack - as discussed in the main text - can be attributed to the way that it handles noising and denoising of the side-chain *χ* angles. We observed that DiffPack uses the ground-truth bond angles and bond lengths without idealization during training and inference ^2^, which is an issue due to two main reasons: 1. inference is not possible when the ground-truth side-chain coordinates are not known (ex. samples from backbone generative models), and 2. *a priori* information of input side-chain conformations should not impact predictive performance. The first reason is a trivial issue since any random initialization of torsion angles can be easily applied to backbone-only structures to serve as a starting point for side-chain packing tools. However, we discovered that for structures where ground-truth bond angles and bond lengths are unknown (i.e. using idealized structures), DiffPack performs notably worse, limiting applicability to side-chain packing for structures without access to ground-truth sidechain conformations.

First, we verify that the ground-truth bond angles and bond lengths are unchanged before and after DiffPack in Supplementary Figure 1. We analyze both Ca-C*β* bond length and N-Ca-C*β* bond angles for DiffPack and FlowPacker, and observe that DiffPack perfectly recapitulates the ground-truth distributions of both bond angles and bond lengths, while FlowPacker does not since it uses idealized bond angles and lengths.

To assess the performance discrepancy of using varying structures that **only differ in the side-chain coordinates**, we input various structures to both DiffPack and AttnPacker, one of the side-chain packing baselines used in this paper, and report the results in Supplementary Table 1. Note that in all the different input structures, the backbone coordinates are unchanged. First, we report that idealization of bond lengths and angles results in a *>* 0.25*A* increase in RMSD, which provides significant advantage to DiffPack given that side-chain packing tools attains sub-Angstrom accuracy. Moreover, we observe that DiffPack shows worse performance when we use idealized, Rosetta-packed, and AttnPacker-packed structures as input, suggesting that the ground-truth bond angles and lengths contribute significantly to DiffPack’s performance. On the other hand, AttnPacker shows the same performance across all inputs, since it is agnostic to the input sidechain coordinates. Therefore, we conclude that DiffPack does exhibit data leakage to some extent, since side-chain packing models should only be dependent on the backbone coordinates and not the input structure’s side-chain coordinates. The current version of DiffPack may be useful when searching sidechain conformational states of known protein structures, but its applicability to structures without knowledge of the ground-truth side-chain conformations may be limited. We also note that DiffPack’s performance may improve when retrained with idealized bond angles and lengths. We have contacted the authors of DiffPack to notify them of our findings.

**Figure S1:**
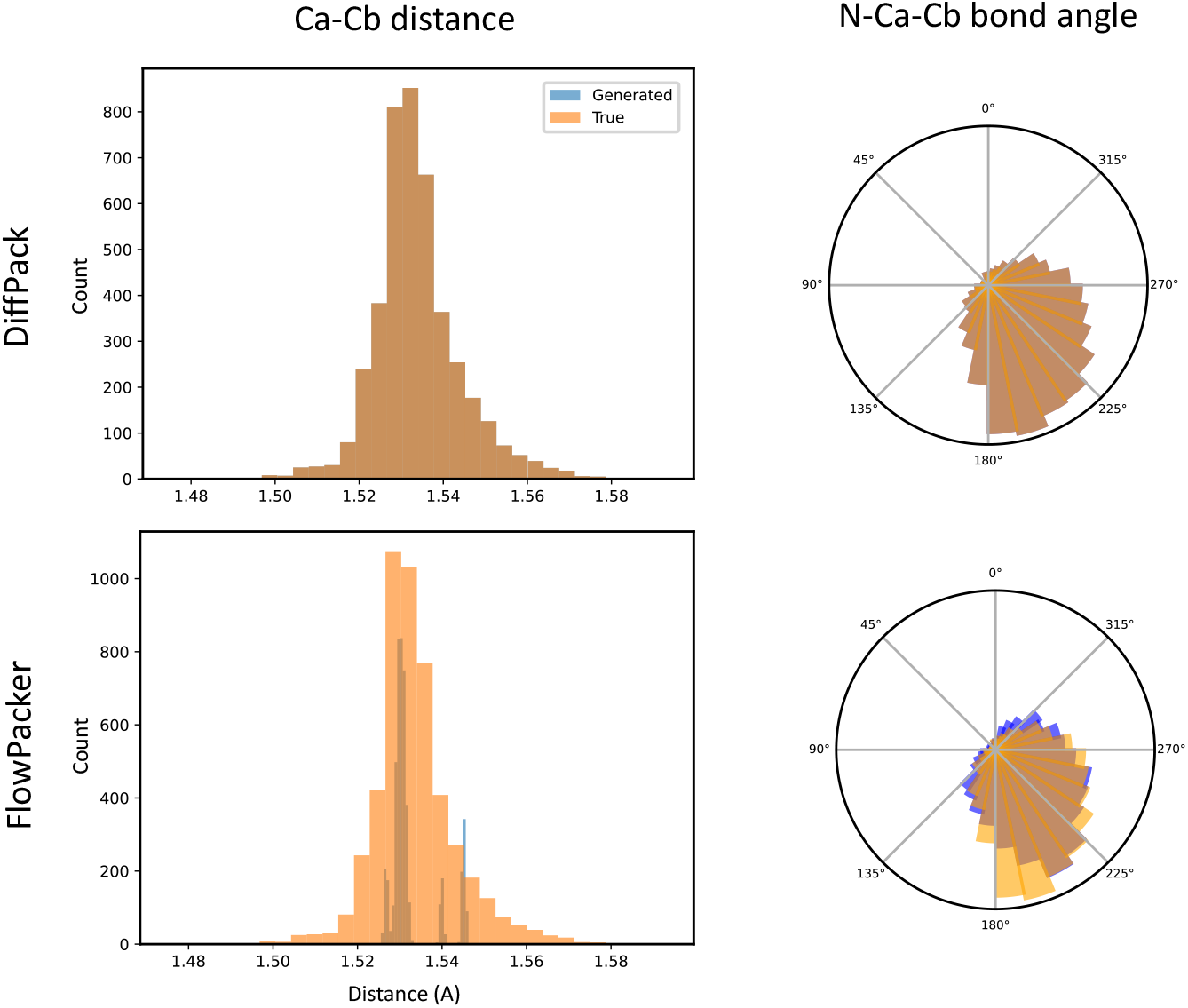
Analysis of N-Ca-C*β* bond angle and Ca-C*β* bond lengths before and after DiffPack and FlowPacker. We observe that DiffPack perfectly recapitulates the input structure’s bond angle and lengths, which we believe may be a source of data leakage since it is not fully agnostic to the ground-truth sidechain. FlowPacker uses idealized coordinates given the noised torsions, which results in different distributions from the input ground-truth structures. Note that the variations in idealized distances of FlowPacker is a result of using ground-truth Ca coordinates with idealized C*β* ones. The plots were generated by using the CASP13 test set with the respective models.

**Table S1:** Performance metrics of DiffPack and AttnPacker on various input structures on the CASP13 test set. We observe that idealization of bond angles and lengths results in a >0.25A RMSD from the ground-truth, suggesting models that directly use the ground-truth bond angles and lengths significantly benefits atom RMSD performance over idealized or agnostic packing tools.

| Model | Input | ANGLE MAE $^{\circ}$ ↓ | | | | RMSD $\text{\AA}$ ↓ |
| --- | --- | --- | --- | --- | --- | --- |
| | | $\chi_1$ | $\chi_2$ | $\chi_3$ | $\chi_4$ | All |
| Idealizer | Ground Truth | 1.33 | 0.02 | 0.03 | 0.03 | 0.254 |
| DiffPack | Ground Truth | 14.07 | 23.20 | 36.79 | 48.13 | 0.571 |
|  | Idealized | 22.26 | 29.96 | 46.61 | 53.47 | 0.778 |
|  | Rosetta | 21.43 | 29.90 | 46.20 | 54.25 | 0.771 |
|  | AttnPacker-pp | 21.98 | 29.22 | 45.35 | 54.77 | 0.780 |
| AttnPacker | Ground Truth | 16.33 | 27.46 | 50.42 | 49.40 | 0.677 |
|  | Idealized | 16.33 | 27.46 | 50.42 | 49.40 | 0.677 |
|  | Rosetta | 16.33 | 27.46 | 50.42 | 49.40 | 0.677 |
|  | AttnPacker-pp | 16.33 | 27.46 | 50.42 | 49.40 | 0.677 |

**Figure S2:**
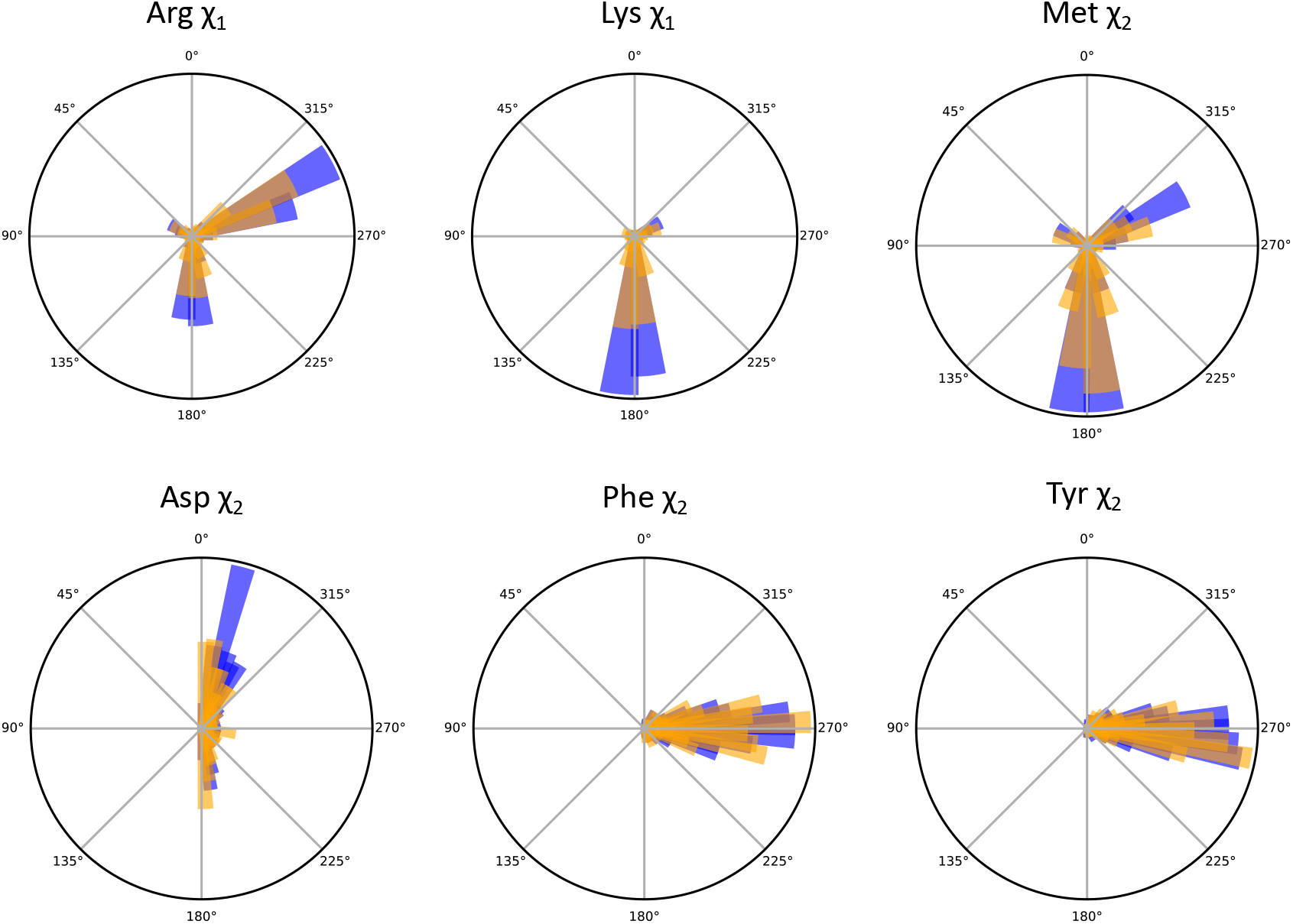
Generated vs. true *χ* distributions. Three randomly selected *χ* distributions are listed on the top row, while the bottom row contains three *π*-symmetric *χ* angles that lie in the interval [0, 180] due to the parameterization in FlowPacker. Generated *χ* angles are depicted in orange, while ground-truth *χ* angles are in blue. The distributions are generated using the CASP13 test set.

**Figure S3:**
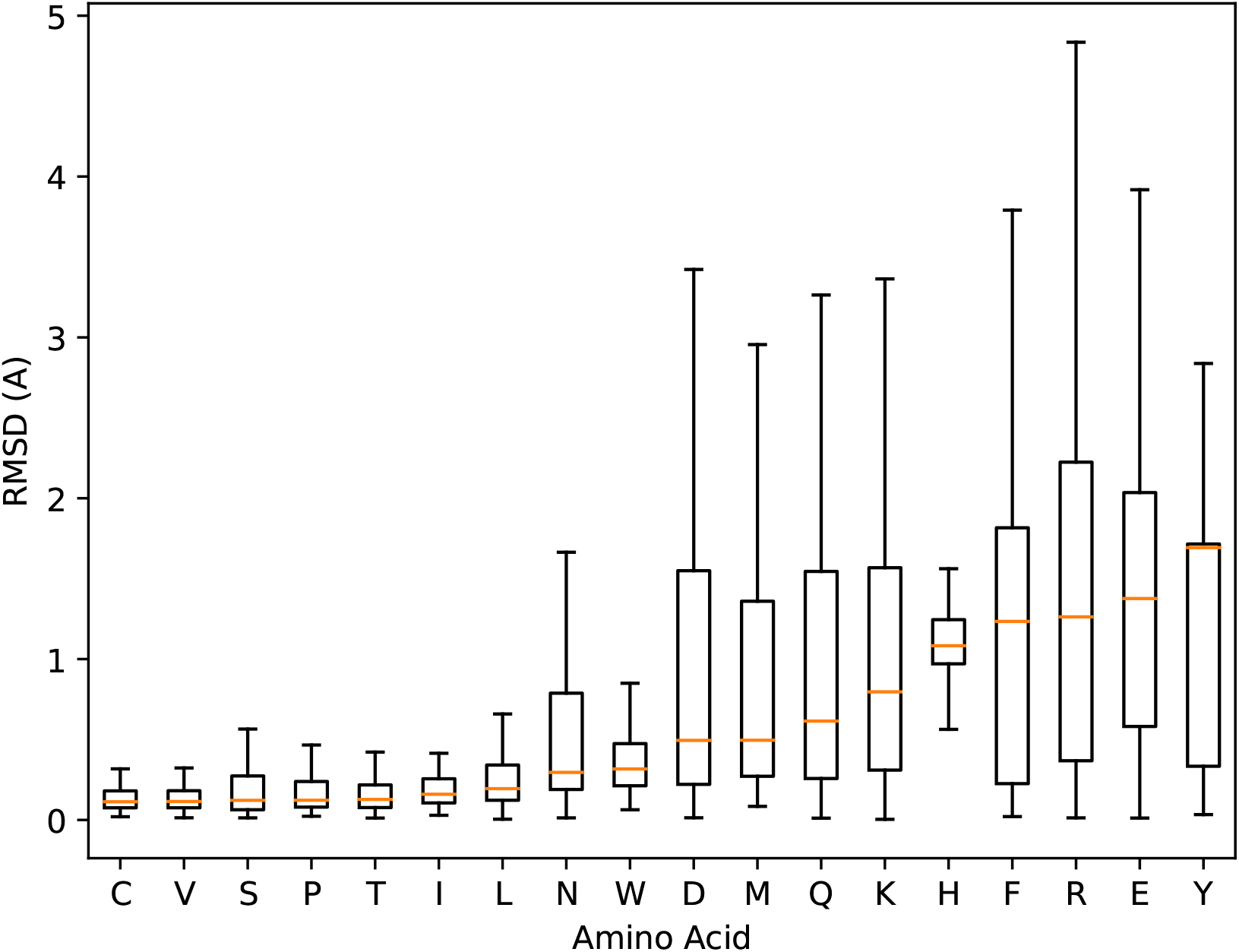
RMSD distributions per amino-acid identity, ranked from lowest to highest median atom RMSD. We observe that aromatic (Y, F, H) and long linear (E, R, K) sidechains exhibit the highest median RMSD.

**Table S2:** Performance evaluation of FlowPacker on CASP15 trained on two different datasets.

| Dataset | ANGLE MAE ° ↓ |  |  |  | ANGLE ACCURACY % ↑ |  |  |  | ATOM RMSD Å ↓ |  |  |
| --- | --- | --- | --- | --- | --- | --- | --- | --- | --- | --- | --- |
| | $\chi_1$ | $\chi_2$ | $\chi_3$ | $\chi_4$ | $\chi_1$ | $\chi_2$ | $\chi_3$ | $\chi_4$ | All | Core | Surface |
| BC40 | 24.82 | 31.14 | 43.74 | 56.68 | 77.14% | 67.46% | **48.90%** | 45.61% | 0.800 | 0.435 | 1.027 |
| PDB-S40 | **23.50** | **29.66** | **43.22** | **54.37** | **78.04%** | **68.71%** | 48.69% | **46.42%** | **0.770** | **0.404** | **0.991** |

## Footnotes

1 Although the need for equivariance has been challenged by recent works, notably AlphaFold3 [2], since equivariant networks often are more expensive to train over optimized Transformer architectures [20].

2 see rotate_side_chain function in https://github.com/DeepGraphLearning/DiffPack/blob/main/diffpack/rotamer.py

